# Evolutionary rate covariation identifies SLC30A9 (ZnT9) as a mitochondrial zinc transporter

**DOI:** 10.1101/2021.04.22.440839

**Authors:** Amanda Kowalczyk, Omotola Gbadamosi, Kathryn Kolor, Jahree Sosa, Claudette St Croix, Gregory Gibson, Maria Chikina, Elias Aizenman, Nathan Clark, Kirill Kiselyov

**Affiliations:** Joint Carnegie Mellon University-University of Pittsburgh PhD Program in Computational Biology, Pittsburgh, USA, 15213; Department of Computational and Systems Biology, University of Pittsburgh, USA, 15213; Department of Biological Science, University of Pittsburgh, USA, 15260; Center for Biologic Imaging, University of Pittsburgh, USA, 15260; Department of Neurobiology and Pittsburgh Institute for Neurodegenerative Diseases, University of Pittsburgh School of Medicine, Pittsburgh, PA 15260; Department of Human Genetics, University of Utah, USA, 84112

## Abstract

Recent advances in genome sequencing have led to the identification of new ion and metabolite transporters, many of which have not been characterized. Due to the variety of subcellular localizations, cargo and transport mechanisms, such characterization is a daunting task, and predictive approaches focused on the functional context of transporters are very much needed. Here we present a case for identifying a transporter localization using evolutionary rate covariation (ERC), a computational approach based on pairwise correlations of amino acid sequence evolutionary rates across the mammalian phylogeny. As a case study, we find that poorly characterized transporter SLC30A9 (ZnT9) uniquely and prominently coevolves with several components of the mitochondrial oxidative phosphorylation chain, suggesting mitochondrial localization. We confirmed this computational finding experimentally using recombinant human SLC30A9. SLC30A9 loss caused zinc mishandling in the mitochondria, suggesting that under normal conditions it acts as a zinc exporter. We therefore propose that ERC can be used to predict the functional context of novel transporters and other poorly characterized proteins.

## Introduction

Localization in a specific organelle or subset of organelles defines the functional context for many molecules, especially ion and metabolite transporters. For example, there are ion channels and transporters that localize to the plasma membrane or endo/sarcoplasmic reticulum, and even though they might conduct the same ions, they clearly have different impact on cellular function (Butorac et al., 2020; Gandini and Zamponi, 2021; Guse et al., 2021; Lemos et al., 2021; Lopez et al., 2020). Similarly, a large cohort of zinc transporters, responsible for pumping ionic zinc into distinct organelles or in or out of cells, subserve distinct functions, including synaptic modulation (Krall et al., 2021) and co-secretion with milk, in the latter case providing an essential dietary supplement and shaping mammary gland development (McCormick et al., 2014). The recent advances in genomic sequencing have resulted in identification of many putative transporters. While some aspects of their function can be inferred based on sequence homology or structural similarity to previously characterized molecules, important details, such as subcellular localization, are usually beyond the reach of these approaches. The main objective of the present studies is to evaluate whether a computational approach based on molecular evolution can accurately predict cellular localization of poorly characterized cellular proteins. Here, this idea was tested using zinc transporters as a model.

An example of ubiquitous but poorly understood transporters are the zinc transporter families. Zinc is an essential component of cellular function, as a key cofactor in enzymatic reactions, transcription and synaptic transmission (Kambe et al., 2015). Moreover, it has been estimated that approximately 10% of human proteome binds zinc (Andreini et al., 2006). Zinc deficit is a factor in growth delays, hair and skin pathologies and behavioral defects (Prasad, 2013). On the other hand, excess zinc promotes the production of reactive oxygen species and many forms of cell damage (Valko et al., 2005). Because of zinc’s duality, cellular zinc concentrations are tightly regulated by a system of transporters and metal-binding proteins. In particular, zinc entry into the cytoplasm is managed by the SLC39A (Zip) family of transporters, while its expulsion or sequestration into organelles is driven by the SLC30A family. Metallothioneins, in addition to other zinc-binding proteins, provide an important intracellular buffer for the metal, leading extremely low (sub nanomolar), or perhaps non-existent, levels of free zinc within the cytoplasm (Thirumoorthy et al., 2011). The zinc transporter families consist of two dozen structurally related members discovered as a result of advances in sequencing technology and gene annotation (Kambe et al., 2015; Schweigel-Röntgen, 2014). However, many of these molecules remain under-characterized at the level of subcellular localization or transport mechanisms. The full extent of annotations supported by several lines of experimental evidence indicate that, for example, SLC30A1 and SLC39A1 function in the plasma membrane (Milon et al., 2006; Palmiter and Findley, 1995), SLC30A3 functions in synaptic vesicles (Palmiter et al., 1996) and SLC30A2 and SLC30A4 function in lysosomes and related organelles (Kukic et al., 2014, 2013; Lopez and Kelleher, 2009). For many of the remaining SLC30A and SLC39A transporters, however, functional annotations, including cellular localizations, are fragmentary. Importantly, some organelles appear to contain several zinc transporters with overlapping functions, and the reason for this is unclear (Kambe et al., 2017, 2015).

At the same time, some key aspects of organellar zinc transport remain poorly understood and/or lack an assigned transporter. This is especially the case for mitochondrial zinc, which, as a cofactor of several mitochondrial enzymes, is a critical component of mitochondrial physiology (Granzotto et al., 2020; Ji et al., 2020; Turan and Tuncay, 2021). In parallel to full-cell consequences of zinc dysregulation, mitochondrial zinc overload is toxic and has been linked to traumatic brain injury and stroke (Frederickson et al., 2004; Galasso and Dyck, 2007; Granzotto et al., 2020; Isaev et al., 2020; Ji et al., 2019; Nolte et al., 2004; Pan et al., 2015; Qi et al., 2019; Sensi et al., 2011, 2009; Yin et al., 2019; Zhao et al., 2018). The mechanisms of mitochondrial zinc regulation are poorly understood and neither SLC30A nor SLC39A members have been directly shown to localize in the mitochondria. Moreover, while the mitochondrial calcium uniporter MCU was proposed to be responsible for zinc uptake into the mitochondria from the cytoplasm (Clausen et al., 2013; Ji et al., 2020; Medvedeva and Weiss, 2014), the pathway that dissipates the mitochondrial zinc has not been described.

These considerations illustrate a need for a predictive approach to match zinc transporters to specific organelles and focus experimental hypotheses to a reasonable set of targeted studies. We propose Evolutionary Rate Covariation (ERC) as the link between functional genomic information about zinc transporters and their associated organelles. ERC is based on the central hypothesis that proteins with shared functional context will share evolutionary pressure and thus evolve at similar rates, even as their rates change over time and between species (Clark et al., 2012; Priedigkeit et al., 2015). This evolutionary rate covariation occurs in physically interacting proteins that coevolve as well as functionally related proteins such as gene regulatory elements, membrane traffic adaptors, and transporters. As a result, ERC has been used to identify the functions of uncharacterized genes, including novel DNA damage-induced apoptosis suppressor, sex peptide network components, regulators of cell adhesion, and ion and aminoacid transporters (Brunette et al., 2019; Clark et al., 2012; Findlay et al., 2014; Raza et al., 2019; Talsness et al., 2020; Ziegler et al., 2016). ERC is calculated as correlation between evolutionary rates throughout a phylogeny. In fact, the work presented here will focus on values calculated across 33 mammal species. Genes are inferred to be functionally related if they have a high degree of correlation in evolutionary rates, or, in other words, if the genes in question have similar evolutionary rates across species.

We present ERC as a powerful tool for de-novo prediction of molecular function with characterization of the zinc transporter SLC30A family, specifically SLC30A9, its most poorly characterized member. Although there are previously reported links to zinc transport (Perez et al., 2017; Zhang et al., 2015), this transporter had previously not been linked to an organellar function in mammalian cells. We show here that ERC analysis suggests a strong and specific signal for SLC30A9 coevolution with several components of the mitochondrial oxidative phosphorylation chain including complex I, and the mitochondrial H^+^ driven ATP synthase (complex V (Jonckheere et al., 2012; Neupane et al., 2019; Sazanov, 2015; Zhao et al., 2019)). The ERC signal between SLC30A9 and mitochondrial complexes was significantly higher than the signal between SLC30A9 and the vacuolar/vesicular/lysosomal H^+^ pump. We also find that recombinant SLC30A9 co-localizes with the mitochondrial protein marker TOM20, strongly suggesting mitochondrial localization, consistent with some prior bioinformatics evidence (Calvo et al., 2016; Rath et al., 2021). Furthermore, SLC30A9 knockdown in HeLa cells using siRNA suppresses the dissipation of mitochondrial zinc after zinc overload. We therefore propose that SLC30A9 is a mitochondrial zinc exporter and thus possesses a unique functional profile unique for members of the SLC30A family. Of interest, further inquiry of SLC30A sequence revealed that SLC30A9 has followed a unique evolutionary trajectory -- it is deeply and highly conserved from mammals through archaea and proteobacteria, while other SLC30As are likely a result of more recent gene duplication events. These findings both illuminate the function of SLC30A9 and distinguish it as a unique molecule among SLC30A family members.

## Results

### SLC30A9 phylogeny

We sought to investigate SLC30A9 function by analyzing its evolution throughout the tree of life. Although SLC30A9 variants have been previously identified in some species, including rat, mouse, zebrafish, and fruit fly (Pubmed search), we performed a more rigorous, targeted search. Due to heterogeneity of the SLC30A/SLC39A superfamily, we began by establishing defining features of SLC30A9 and other SLC30As. Zinc permeation by SLC30A is coordinated by aspartate and histidine residues contributed by the HxxxD motif (flanked by D at position 39 of human SLC30A1, forming DxxHxxxD signature) and by the HxxxD motif located downstream of the first motif (at position 250 of human SLC30A1) (**Fig 1A; full alignment in Fig S1**). Previously published crystal structures and homology modeling show that these motifs are located in the transmembrane domains 2 and 5 of human SLC30As (Fukada and Kambe, 2011; Hoch et al., 2020, 2012; Ohana et al., 2009). Amino acid alignment of human and other organisms’ SLC30A9 with other SLC30As, coupled with homology modeling, reveals a distinct and conserved arrangement (**Fig 1A and S1**). In human SLC30A9, the HxxxD sequence is flanked by a glutamate residue (ExxHxxxD at position 268 of human SLC30A9) and the downstream sequence is ILLED.

These substitutions provide a unique signature that identifies SLC30A9 and distinguishes it from other SLC30A family members. The signatures are surprisingly well conserved: homologous sequences containing these substitutions and transmembrane domain arrangements reminiscent of human SLC30A9, are found throughout the tree of life (**Fig 1A and S1**). Critically, a detailed phylogenetic analysis of SLC30A9 candidates and other members of the SLC30A family identifies SLC30A9 as a distinct phylogenetic entity, components of which are more related to each other across the tree of life, than to other SLC30A members.

**Figure 1:**
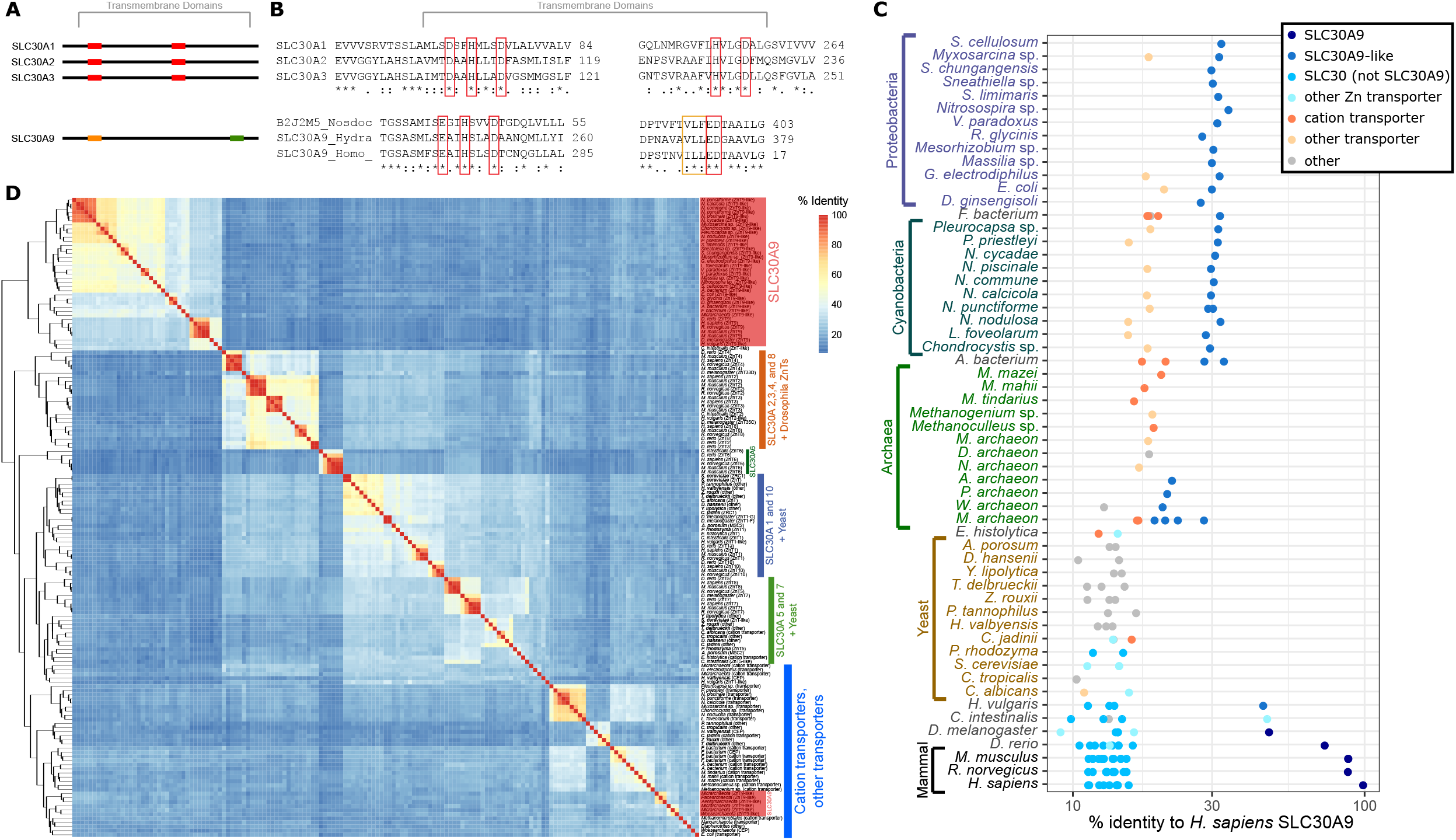
Defining features of SLC30A9. A) Cartoon indicating defining features of SLC30A9 and other SLC30As. B). Partial amino acid alignment of select human SLC30A (top) and SLC30A9 from humans and other organisms (bottom), specifically focused on the conserved differences in the putative permeation center depicted in part A. C) Bacterial SLC30A9s show higher percent identity to human SLC30A9 than do other animal SLC30As. This suggests potential divergence of SLC30A9 before animal duplication events give rise to other SLC30As. D) Hierarchical clustering based on percent sequence identity demonstrates that bacterial and animal SLC30A9s cluster together separately from other animal SLC30As, yeast transporters, cation transporters, and other ion transporters, further supporting ancient divergence of SLC30A9 prior to duplication of other SLC30As.

The substitutions specific to SLC30A9 appear to be exceptionally well-conserved as we can detect SLC30A9 homologous sequences containing the invariant ExxHxxxD…(I/V/L)(I/V/L)xED in every clade except for the yeast and other fungi (**Table 1, Fig 2A**). This includes archaebacteria, proteobacteria, cyanobacteria, firmicutes, and actinobacteria, as well as every animal species tested. Organisms on which SLC30A9 is present typically contain a single SLC30A9 homolog, with the exception of *Nosdoc punctiforme* (2 copies), *Actinobacteria bacterium* (2 copies), and *Micrarchaeota archaeon* (3 copies). The SLC30A superfamily increases in complexity in higher organisms, with ten members (including SLC30A9) in mammals, 6 members in *Ciona intestinalis* and *Drosophila melanogaster*, 4 members in *Hydra vulgaris*, and typically 1 or 2 copies in most bacteria and some yeast species. However, of important note, the numerous duplications of SLC30As are not paralleled by increasing numbers of SLC30A9 homologs in higher-order organisms (**Fig 2A**). This distinction points toward a unique evolutionary trajectory for SLC30A9 compared to other SLC30s, potentially because of a unique molecular function for this transporter.

**Table 1 legend.** Expression of SLC30A9, MCU, MICU, MMT and their homologs in various organisms (x shows expression).

| Clade | Species | SLC30A9 | MCU | MICU | MMT |
| --- | --- | --- | --- | --- | --- |
| **Animals** | Homo sapiens | x | x | x |  |
|  | Mus musculus | x | x | x |  |
|  | Rattus norvegicus | x | x | x |  |
|  | Danio rerio | x | x | x |  |
|  | Ciona intestinalis | x | x | x |  |
|  | Hydra vulgaris | x | x | x |  |
|  | Drosophila melanogaster | x | x | x |  |
| **Fungi** | Yarrowia lipolytica |  |  |  | x |
|  | Saccharomyces cerevisiae |  |  |  | x |
|  | Debaryomyces hansenii |  |  |  | x |
|  | Pachysolen tannophilus |  |  |  | x |
|  | Williopsis saturnus |  |  |  |  |
|  | Ustilago maydis 521 |  | x |  | x |
|  | Cryptococcus neoformans |  | x |  | x |
|  | Laccaria bicolor S238N-H82 |  |  |  | x |
|  | Magnaporthe oryzae 70-15 |  | x |  | x |
|  | Penicillium chrysogenum 54-1255 |  | x |  | x |
|  | Uncinocarpus reesii 1704 |  |  |  | x |
| **Algae** | Ostreococcus lucimarinus CCE9901 | x | x | x |  |
|  | Chlamydomonas reinhardtii | x | x | x |  |
|  | Volvox carteri f. nagarensis | x | x | x |  |
| **Amoeba** | Pelomyxa palustris |  |  |  |  |
|  | Chaos carolinense |  |  |  |  |
|  | Amoeba proteus |  |  |  |  |
|  | Entamoeba histolytica |  |  |  |  |
|  | Arcella vulgaris |  |  |  |  |
|  | Vitis vinifera | x | x | x |  |
|  | Populus trichocarpa | x | x | x |  |
|  | Lotus japonicus |  | x | x |  |
| **Protists** | Trichomonas vaginalis G3 |  |  |  |  |
|  | Leishmania infantum JPCM5 |  | x | x |  |
|  | Trypanosoma brucei TREU927 |  | x | x |  |
|  | Tetrahymena thermophila SB210 |  | x | x |  |
|  | Paramecium tetraurelia strain d4-2 |  | x |  |  |
|  | Phaeodactylum tricornutum CCAP 1055/1 | x |  |  |  |
|  | Plasmodium chabaudi chabaudi |  |  |  |  |
| **Archaea** | Candidatus Pacearchaeota archaeon | x |  |  |  |
|  | Candidatus Woesearchaeota archaeon | x |  |  |  |
|  | Candidatus Aenigmarchaeota archaeon ex4484_52 | x |  |  |  |
|  | Candidatus Diapherotrites archaeon |  |  |  |  |
|  | Methanohalophilus mahii |  |  |  |  |
|  | Methanobrevibacter smithii |  |  |  |  |
|  | Methanosarcina mazei |  |  |  |  |
| **Bacteria** | Sneathiella chungangensis | x |  |  |  |
|  | Massilia sp. LC238 | x |  |  |  |
|  | Geopsychrobacter electrodiphilus | x |  |  |  |
|  | Escherichia Coli | x |  |  |  |
|  | Chondrocystis sp. NIES-4102 | x |  |  |  |
|  | Nosdoc punctiforme | x |  |  |  |
|  | Leptolyngbya foveolarum | x |  |  |  |
|  | Nostoc cycadae | x |  |  |  |

**Figure 2.**
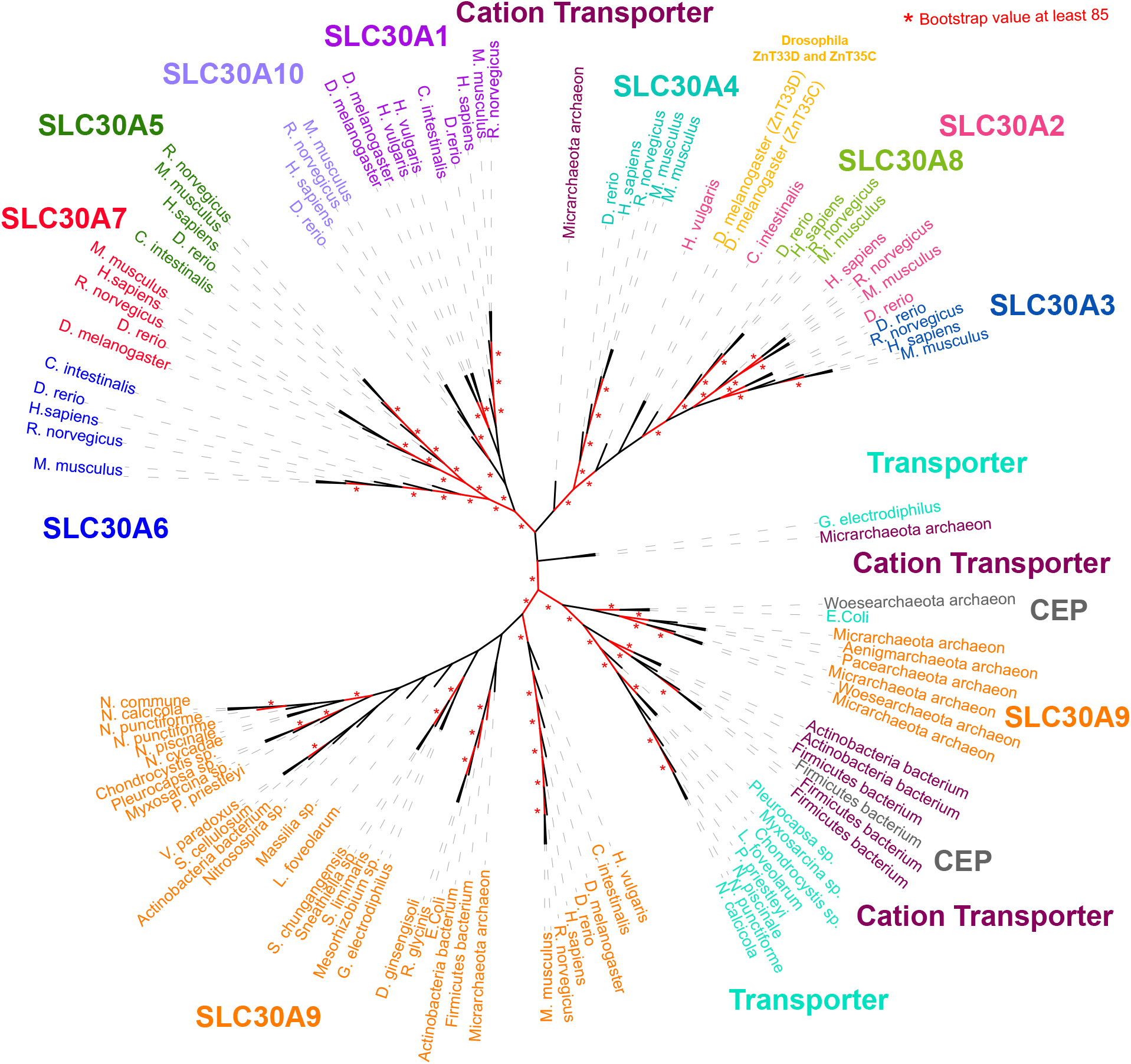
SLC30A dendrogram suggests ancient divergence of SLC30A9 and other zinc transporters. The branch leading to all SLC30A9s, both mammalian and bacterial, as well as some other transporters, is supported with high bootstrap confidence. Other SLC30As in animals and some other transporters branch separately.

As further evidence of SLC30A9’s unique evolution, the human SLC30A9 is more closely related to bacterial SLC30A9 than to other human SLC30As based both on sequence identity and on clustering through sequence-based phylogenetic tree construction (**Fig 2B**). The SLC30A9 sequences appear to be an ancient clade and have remained a single-copy orthologous group during eukaryotic and bacterial evolution. This monophyletic relationship of the ancient SLC30A9 subtree is supported by strong branch support (100% bootstrap support, indicating high fidelity of placing SLC30A9 as an independent tree branch). The phylogenetic tree also shows a clade of SLC30A9 sequences from Archaea that may not be monophyletic with the rest of the SLC30A9 clade, making their membership in this family uncertain. Together, these findings show both that SLC30A9 is an ancient, deeply conserved protein, and that it could be considered apart from the rest of the eukaryotic SLC30A family. As such, it appears that the ancestor of the SLC30A family was present early in Prokaryote evolution. Indeed, SLC30A9 has remained relatively static, not undergoing further duplications. In contrast, the remaining SLC30A family members have gone through multiple gene duplications and deletions, leading to the SLC30A1-8 and 10 paralogs in Eukaryotes.

Yeast and other fungi do not appear to have SLC30A9. MMT1/2 yeast transporters, which have been suggested to be the yeast analog of SLC30A9, do not have the ExxHxxxD…(I/V/L)(I/V/L)ED substitution, and thus likely have a different function. Nonetheless, there appears to be mutual exclusivity between SLC30A9 and MMT1/2 expression, as organisms containing SLC30A9 do not seem to have MMT1/2 (**Table 1, Fig 2A**), suggesting that MMT1/2 may replace SLC30A9 in those species, although our sequence identity and phylogeny construction analyses suggest that MMT1/2 would not have originated from substitutions in SLC30A9. Although we did not further pursue this avenue of inquiry, mechanisms by which yeast species compensated for the loss of such a deeply conserved protein is of great interest. MMT1/2 are found in the yeast mitochondria. This, together with the correlation between the presence of SLC30A9 and MCU in many species (**Table 1**) strongly suggested that SLC30A9 would be present in the mitochondria.

### SLC30A9 coevolution with the mitochondrial components

Since we found that SLC30A9 is an evolutionarily distinct and ancient protein, we next applied another evolutionary method to generate a hypothesis about its function and localization. To gain an insight into the cellular localization of SLC30A9 we performed an ERC analysis between SLC30A9 and a battery of other organellar transporters. Because some SLC30A transporters have been shown or proposed to use a proton gradient as a driving force for zinc transport (Golan et al., 2019; Hoch et al., 2020; Ohana et al., 2009), our rationale for this evolutionary approach was as follows: if SLC30A9 uses an ionic gradient to pump zinc, then its evolution could be influenced by the same evolutionary pressures acting on the transporter that establishes the ionic gradient. Hence, the two transporters will have correlated evolutionary rates, which will be uncovered by ERC. We can, therefore, use well established ion transporters, whose localization is known, as baits to search for SLC30As that have coevolved with them, and, as such, probably located within the same organelle.

The analysis was performed using a publicly available ERC portal (https://csb.pitt.edu/erc_analysis/) (Wolfe and Clark, 2015). This approach utilizes gene-specific evolutionary rates from 33 mammalian species and their ancestral branches calculated for more than 19,000 protein-coding genes. A gene pair’s ERC value is calculated as the correlation coefficient between their rates in those species. Analysis of SLC30A9 focused on several organellar transporters including the vacuolar/vesicular/lysosomal H^+^ pump (multimolecular complex comprising structurally unrelated proteins that are coded by the ATP6 family of genes (Beyenbach and Wieczorek, 2006; Grabe et al., 2000)), the endolysosomal ClC transporters (Jentsch, 2007; Poroca et al., 2017), and the components of the mitochondrial oxidative phosphorylation chain (Chaban et al., 2014; Kühlbrandt, 2015; Lippe et al., 2019; Sousa et al., 2018), which either establish or utilize the mitochondrial H^+^ gradient (see Materials and Methods and **Fig S2** for the list). We calculated ERC values between SLC30A9 and each of the genes annotated to those batteries of transporters. MCU coevolution with MICU1 (Fan et al., 2020; Mallilankaraman et al., 2012; Patron et al., 2014) was used as a positive control for coevolution of functionally linked proteins (R value 0.584, p=0.002).

SLC30A3 and SLC30A4, known to localize in the acidic organelles containing high levels of ATP6, namely synaptic vesicles (Cole et al., 1999; Palmiter et al., 1996) and lysosomes (Kukic et al., 2013), respectively, were used as controls for specificity of the observed effects. Both of these transporters were shown, or proposed, to utilize H^+^ gradient established by ATP6 pumps in order to move zinc. As predicted in this control analysis, SLC30A3 and SLC30A4 show several significant ERC values with the ATP6 components (**Fig 3A, top, red points**), which is consistent with their cellular localization and H^+^ requirements for transport (Lee and Koh, 2021). In contrast, their ERC scores with the mitochondrial components (blue points) were lower, and, in both cases ERC was skewed towards the vacuolar/vesicular/lysosomal H^+^ pump relative to the mitochondrial signal (**Fig 3A, bottom**).

**Figure 3.**
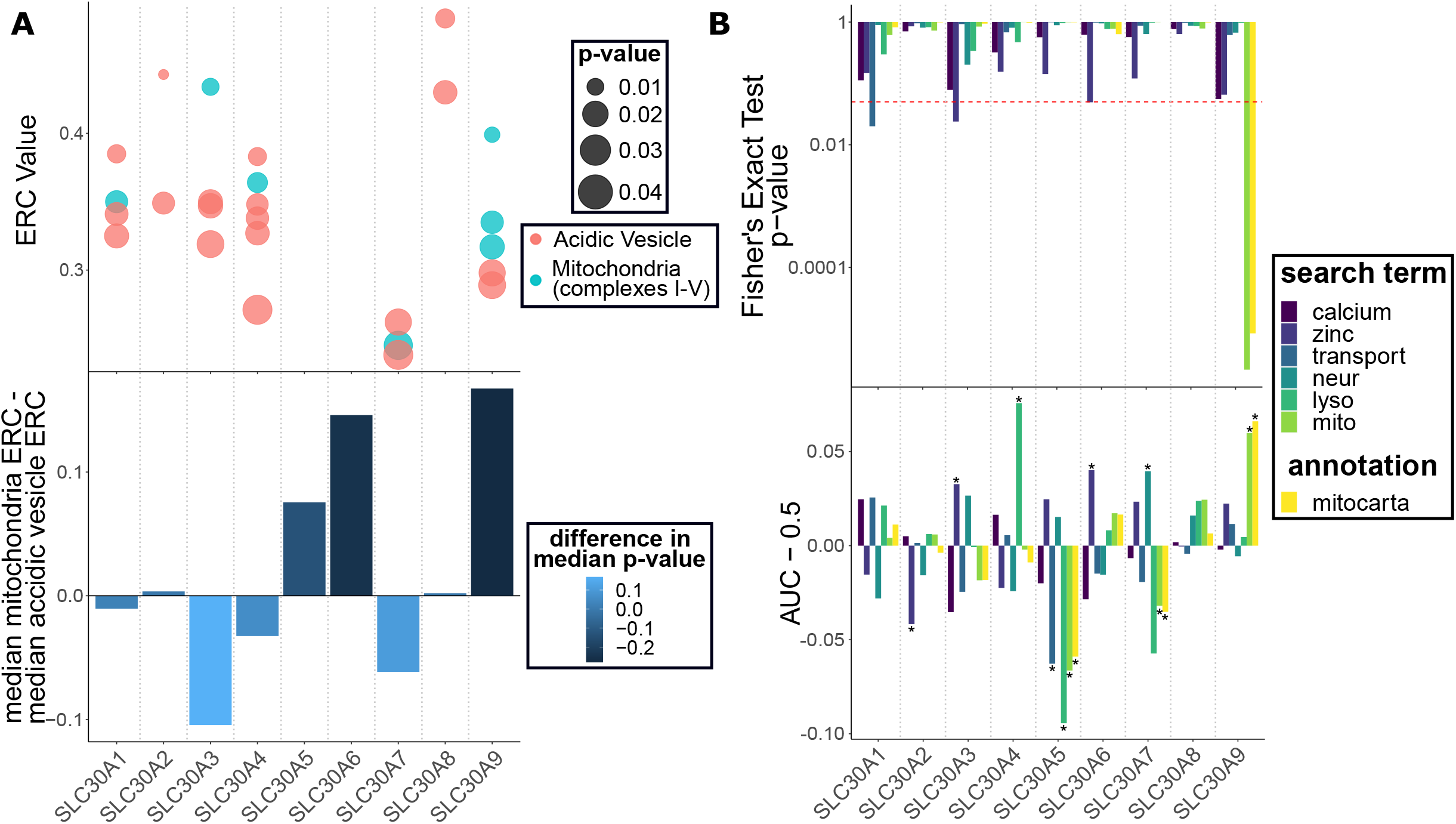
ERC analysis of SLC30 proteins reveals that SLC30A9 (ZnT9) shows unique patterns of correlated evolutionary rates with mitochondrial proteins. A) ERC between SLC30A9 and mitochondrial proteins shows higher correlation and lower p-value compared to ERC between SLC30A9 and lysosomal proteins. This contrast is stronger for SLC30A9 than for any other SLC30. Dots represent genes with positive ERC values and p-values less than 0.05. Bars heights show difference in median ERC value between mitochondrial and lysosomal proteins for all genes and bar colors show difference in median p-value between mitochondrial and lysosomal proteins for all genes. B) SLC30A9 genes with high ERC values are uniquely enriched for mitochondrial function. Search terms represent words identified in gene descriptions to assign them to that term. Also included are Mitocarta annotations. One-tailed Fisher’s Exact Tests were performed to test for significant enrichment of search terms and annotations with ERC values above 0.05, and notably only SLC30A9 shows a significant p-value for mitochondria (bar beyond the red dashed line). A Wilcoxon rank-sum test also showed a uniquely significant positive shift of ERC values for mitochondrial terms and SLC30A9 (asterisks show significance). Centered AUC values demonstrate whether the term was enriched (positive values) or depleted (negative values).

In contrast, we found that SLC30A9 shows a strong coevolutionary signal with several components of the mitochondrial oxidative phosphorylation chain, which is a very unique pattern when compared to other members of the SLC30A family (**Fig 3A, top**). Specifically, it has significantly correlated rates with 3 individual mitochondrial components, belonging to mitochondrial ATP synthase and complex I. In addition, the full distribution of SLC30A9’s scores with the mitochondrial complex encoding genes was much higher than its scores with lysosomal components (**Fig 3A, bottom**).

We next examined the full set of scores for each SLC30A in mammalian genomes to learn with which functions high-scoring genes were associated with it. This study also shows that SLC30A9 is uniquely enriched for high ERC values with genes encoding proteins that are associated with the mitochondria (**Fig 3B**). First, we searched the human RefSeq gene descriptions of the high-scoring genes (ERC value > 0.3) for specific search terms related to calcium, zinc, lysosomes, and mitochondria (Pruitt and Maglott, 2001). SLC30A9’s high-scoring genes were strongly associated with the ‘mito’ search term and with genes annotated as mitochondrial by Mitocarta 2.0 (Fig 3B, upper) (Calvo et al., 2016; Rath et al., 2021). Similarly, we studied the entire distribution of genes as ranked by ERC values with SLC30As. That distribution for SLC30A9 was significantly enriched at high ERC scores for genes with mitochondrial annotations (RefSeq gene description and Mitocarta status) (Fig 3B, lower). In fact, out of all SLC30As, only SLC30A9’s ERC scores showed such associations with mitochondrial annotations.

We also examined the networks of connections drawn based on ERC scores between SLC30As and components of mitochondrial oxidative phosphorylation complexes (**Fig 4A**, upper) and organellar ion transporters, including the vesicular ClC chloride/proton transporters (**Fig 4A**, lower). See also **Fig S2** for more detail. It is clear that SLC30A9 uniquely co-evolves with many mitochondrial components, involving several connections to them, when compared with other SLC30As, as well as other organellar markers. The models in **Fig 4B** summarize the findings. Based on the totality of these evolutionary studies, we propose that SLC30A9 localizes to the mitochondria. We set out next to test this hypothesis.

**Figure 4.**
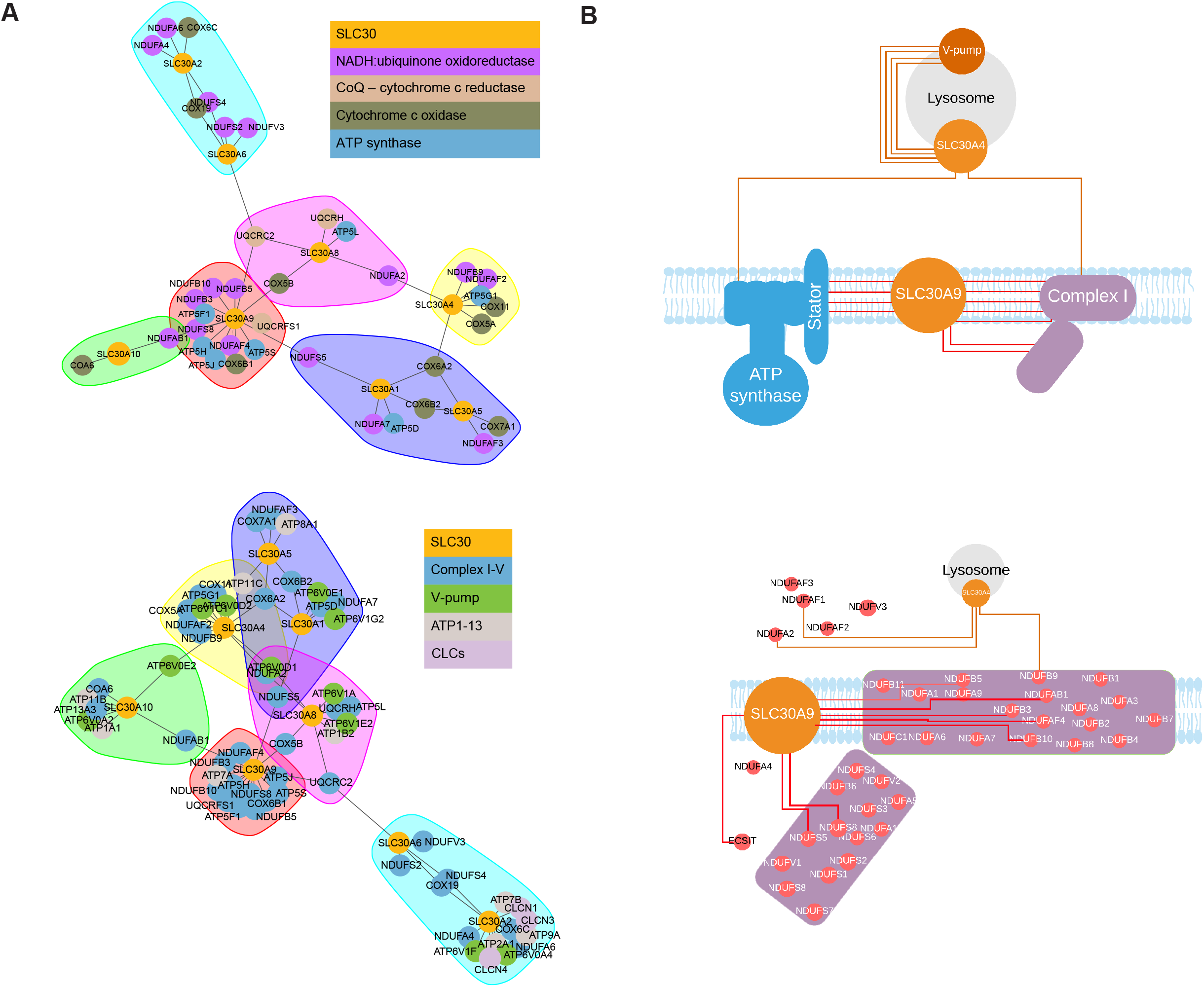
SLC30 evolutionary network clusters and diagrams of SLC30A9 interaction with mitochondrial components. A) Top: SLC30A9 shows robust clustering with numerous mitochondria-related proteins, including NDUF and ATP proteins. Note the cluster including SLC30A9 (in red) with many NADH proteins (purple dots) and ATP synthase proteins (blue dots). Bottom: Clustering using an expanded geneset shows distinctive clustering of SLC30A9 (red cluster) with numerous mitochondria complex I-V proteins (blue circles). B) Wire diagram comparing ERC-positive interactions (at R>0.3) between SLC30A9 and the mitochondrial OxPhos components. SLC30A4 ios shown as a control.

### SLC30A9 localization and effect on mitochondrial zinc

Although the previous bioinformatics assays predict the mitochondrial localization of SLC30A9 (Calvo et al., 2016; Rath et al., 2021), it had not been directly demonstrated. We did not find commercially available antibodies that produce a convincing organellar stain. Therefore, we transfected HeLa cells with recombinant cDNA constructs coding for human SLC30A9 with in-frame C- or N-terminal YFP fusions. **Fig 5** shows that both constructs produced a reticular stain overlapping with the mitochondrial marker TOM20 (Abe et al., 2000). The stretch of positively charged amino acids in the extreme N-terminus of human SLC30A9 is indicative of mitochondrial localization signal, which is supported by Mitoprot II prediction (Claros and Vincens, 1996). Deleting these sequences resulted in significant cell toxicity following transfection, indicating that the mitochondrial localization of SLC30A9 is essential (not shown). Based on this evidence, we conclude that SLC30A9 localizes in the mitochondria. This is the first direct evidence of SLC30A9 localization to this or any other organelle.

**Figure 5.**
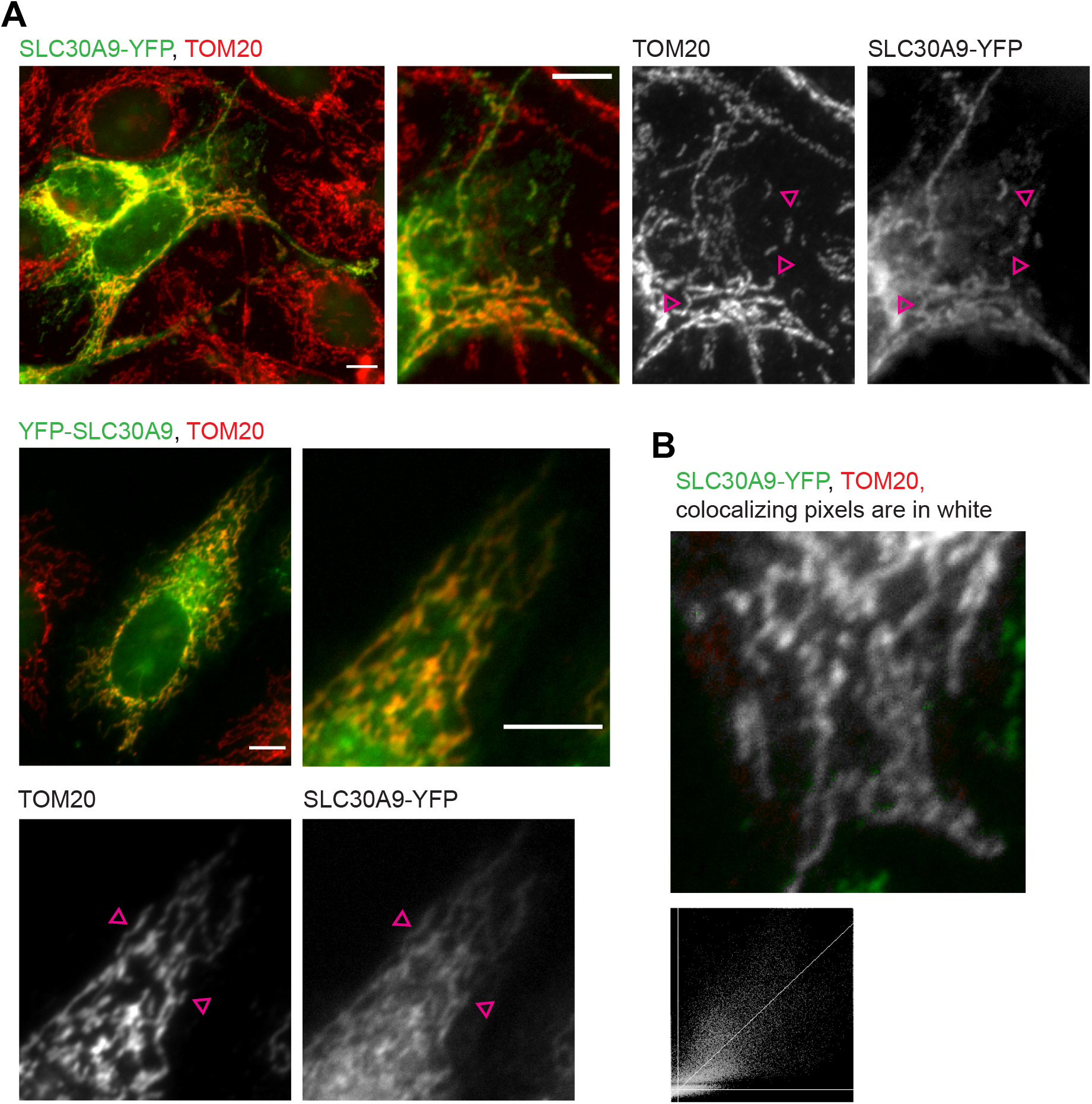
SLC30A9 localization. A. Widefiend fluorescence image of N-terminal (top) and C-terminal (bottom) YFP fusion of human SLC30A9 (green) expressed in HeLa cells and co-stained with antibodies against TOM20 (red). In addition, zoom-in images are shown for clarity. Pink arrows point to examples of overlapping stains. Scale bars are 10 μm. B: An example of colocalization analysis of SLC30A9 and TOM20 obtained using the Colocalization threshold Plugin of FIJI. Histogram below shows linear regression of intensities of the overlapping pixels in the red and green channels.

siRNA-driven SLC30A9 deletion in HeLa cells produced significant changes in mitochondrial zinc handling. Mitochondrial zinc content was analyzed using fluorescent divalent-cation sensitive dye Rhod-2,am, which was loaded into the cells for 15 min, followed by a washout. Fluorescence intensity was analyzed using time-lapse fluorescent imaging; the intensity was measured in the regions of interest which were drawn manually. In zinc-loaded cells, the pattern of the fluorescent signals seemed to conform to the mitochondrial shape. When exposed to 300 μM zinc overnight, SLC30A9 siRNA-transfected cells (knockdown confirmed by qPCR, **Fig 6A**) accumulated significantly more zinc, which is evident from statistically higher fluorescence in these cells, compared with control-transfected cells (**Fig 6B**). Next, we measured the dynamics of the mitochondrial zinc dissipation by returning zinc-exposed cells to a nominally zinc-free medium. Under such conditions, SLC30A9 siRNA-transfected cells lost mitochondria fluorescence signal at a significantly slower rate, indicating slower dissipation of zinc (**Fig 6C**). These data suggest that SLC30A9 is a functional mitochondrial zinc transporter, which, under normal conditions is responsible for zinc extrusion from the mitochondria.

**Figure 6.**
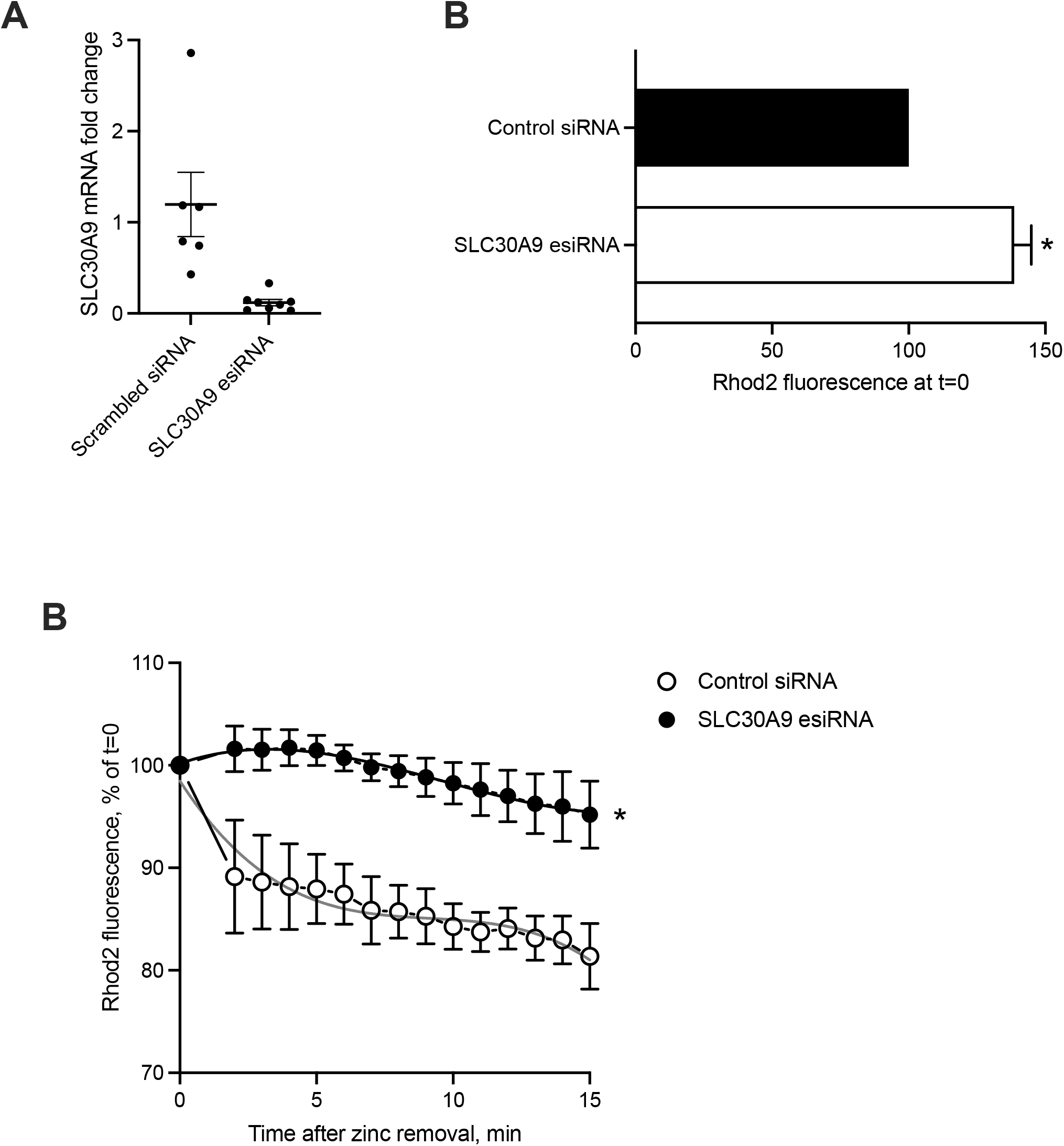
Mitochondrial zinc handling deficits in SLC30A9-knockdown HeLa cells. A) qPCR analysis of HeLa cells transfected with SLC30A9 esiRNA and harvested 24 hours post-transfection. B) Mitochondrial zinc content in control and SLC30A9-transfected HeLa cells was evaluated using Rhod-2,am. The cells were exposed to zinc for 24 hours. Average fluorescence in each region of interest containing clearly identifiable mitochondria was recorded and normalized to the values in the cells that were transfected with scrambled shRNA. C) The delayed loss of mitochondrial zinc in SLC30A9-deficient cells exposed to nominally zinc-tree medium after 24-hour long zinc load. The cells were treated as above; at t=0 the zinc-containing medium was replaced with nominally zinc free medium and images were taken every 60 sec. The fluorescence intensity was normalized to the values recorded at t=0, which was taken as 100%. In Panels A and B, data represent three independent trials involving 7-10 cells. Data are shown ±sem. Asterisks represent statistical significance at p<0.05, for Student’s test in Panel A. In Panel B, the curves were fitted using a third order polynomial curve (smooth gray lines) and an extra sum-of-squares test was performed to answer whether one curve can adequately fit both datasets. That hypothesis was rejected at p<0.0001 level. Here and below, statistical analysis and data plotting were performed using Prism 9.

## Discussion

In the course of the present studies, we used ERC, a bioinformatics assay, to predict the localization of a poorly understood ion transporter. While ERC has been described and tested before (Brunette et al., 2019; Clark et al., 2012; Findlay et al., 2014; Raza et al., 2019; Talsness et al., 2020; Ziegler et al., 2016), our approach utilized a novel idea that tracking evolutionary histories of functionally linked proteins may inform their functional context, such as localization. Therefore, using this approach we were able to predict protein localization in the absence of obvious clearly defined functional domains. We suggest that this approach can have broad applications in diverse biological fields.

Little is known about the SLC30A9 function, although it has recently been linked to Covid-19 because of strong co-purification of SLC30A9 with SARS-CoV-2 proteins (supplementary data in (Gordon et al., 2020)). Furthermore, SLC30A9 has recently been identified as a master regulator of Parkinson’s pathogenesis (Vargas et al., 2021). Given the central role of mitochondria in Parkinson’s pathology (Trinh et al., 2021) and the emerging role of zinc in this process (Park et al., 2014), the evidence of SLC30A9 in the mitochondria suggests a novel contributing factor to Parkinson’s disease.

While SLC30A9 is considered to be a member of the zinc family of transporters, the evidence of its involvement in zinc handling is scarce. Interestingly, polymorphisms in SLC30A9 in African populations have been linked to environmental zinc availability (Zhang et al., 2015), and dysregulation of zinc handling has been detected in fibroblasts from human patients a novel autosomal recessive cerebro-renal syndrome, which has been linked to mutations in *SLC30A9* (Perez et al., 2017). Our data suggest that under normal conditions, SLC30A9 is a mitochondrial zinc exporter. While this seems to be counteractive to the known function of other members of the SLC30A family, namely, moving zinc *into* organelles, it is important to note that the mitochondrial proton gradient operates in the opposite direction in mitochondria, when compared to other membranous organelles such as lysosomes and the Golgi apparatus. Given that some zinc transporters have been shown or proposed to be proton-driven, the “reversed” mitochondrial proton gradient may explain the outward direction of SLC30A9-driven cation transport detected in our system and attributed to SLC30A9 activity.

Mitochondrial zinc transport mechanisms are poorly understood. Several components of the mitochondrial respiratory chain require zinc (Kmita et al., 2015; Suh et al., 2007). Furthermore, proteostasis, protein insertion, enzymatic activities and other functions of the mitochondria are regulated by zinc (Atkinson et al., 2010; Audano et al., 2018; Brambley et al., 2019; Costello et al., 1997; Lee, 2018; Ye et al., 2001). With this in mind, the SLC30A9 coevolution with the mitochondrial oxidative phosphorylation chain components is informative. However, excess zinc has adverse impacts on the mitochondrial respiratory chain and has been implicated in several pathologies including Alzheimer’s disease and stroke (Cherny et al., 2001; Granzotto et al., 2020; Isaev et al., 2020; Lee et al., 2012; Nolte et al., 2004; Pan et al., 2015; Sensi et al., 2009; Wang et al., 2010; Yin et al., 2019; Zhao et al., 2018). Importantly, a mechanism of zinc efflux from the mitochondria had, heretofore, not been identified. Ascribing this role to SLC30A9 is likely to have far-reaching implications in a more complete understanding of management of zinc by the mitochondrial.

At the moment, the mechanistic reasons for coevolution between SLC30A9 and mitochondrial oxidative phosphorylation chain components are not completely clear. ERC suggests functional, but not necessarily physical interaction between these proteins. The coevolution may reflect the shared function of zinc flux and utilization in the mitochondria, as discussed above. Furthermore, SLC30A9 appears to specifically coevolve with the components of the peripheral stalk of the mitochondrial ATP synthase (Fig 4B) suggesting a shared role in proton transport, as the membrane components of the peripheral stalk were suggested to contribute to the formation of the proton channel (Colina-Tenorio et al., 2018). Further investigation would help uncover the mechanism behind the coevolution between SLC30A9 and mitochondrial respiratory chain components and further validate ERC as a cutting-edge tool of molecular discovery and characterization.

## Materials and Methods

### Discovery and categorization of SLC30 sequences

Mammalian and higher eucariote amino acid sequences of SLC30 family members were obtained using Pubmed search and aligned using CLUSTALW (Madeira et al., 2019). The ExxHxxxD…(IVL)(IVL)xED and DxxHxxxD…HxxxD tempates were used as wildcard searches of lower eucaryotes and procaryotes by means of pBLAST. The resulting sequences were verified for the presence of 6 full or partial transmembrane domains using TMHMM server.

### Phylogenetic Analysis

SLC30 family proteins and related homologs collected in the previous step were analyzed in terms of their divergence and phylogeny. Amino acid sequences were aligned using CLUSTALW (Chenna et al., 2003). The resulting alignment was used in PhyML 3.0 to infer their phylogenetic relationships and perform branch support analysis using 100 bootstrap replicates (Guindon et al., 2010). The ‘LG’ amino acid substitution model was used and rate heterogeneity was modeled using a class of invariable sites with freely estimated size and a Gamma rate shape parameter discretized as 4 rate categories. The tree figure in Figure 1 was made using Interactive Tree of Life (iTOL) (Letunic and Bork, 2019, 2007).

### Evolutionary Rate Covariation

The calculation of ERC values was performed as in previous publications (Clark et al., 2012; Priedigkeit et al., 2015). In this case orthologous gene sequences from 33 mammalian species were obtained from the 100-way alignment at the University of California Santa Cruz Genome Browser (Kent et al., 2002). The species chosen were: *Homo sapiens* (human), *Pongo pygmaeus abelii* (orang-utan), *Macaca mulatta* (rhesus macaque), *Callithrix jacchus* (marmoset), *Tarsius syrichta* (tarsier), *Microcebus murinus* (mouse lemur), *Otolemur garnettii* (bushbaby), *Tupaia belangeri* (tree shrew), *Cavia porcellus* (guinea pig), *Dipodomys ordii* (kangaroo rat), *Mus musculus* (mouse), *Rattus norvegicus* (rat), *Spermophilus tridecemlineatus* (squirrel), *Oryctolagus cuniculus* (rab-bit), *Ochotona princeps* (pika), *Vicugna pacos* (alpaca), *Sorex araneus* (shrew), *Bos taurus* (cow), *Tursiops truncatus* (dolphin), *Pteropus vampyrus* (megabat), *Myotis lucifugus* (micro-bat), *Erinaceus europaeus* (hedgehog), *Equus caballus* (horse), *Canis lupus familiaris* (dog), *Felis catus* (cat), *Choloepus hoffmanni* (sloth), *Echinops telfairi* (tenrec), *Loxodonta africana* (elephant), *Procavia capensis* (rock hyrax), *Dasypus novemcinctus* (armadillo), *Monodelphis domestica* (opossum), *Macropus eugenii* (wallaby), and *Ornithorhynchus anatinus* (platypus). Those 17,486 coding sequence alignments were used to calculate gene-specific branch lengths over the mammalian species tree topology using *codeml* of the PAML package (Yang, 2007). For each orthologous gene group/tree, the branch lengths were normalized into relative evolutionary rates through a projection operator (Sato et al., 2005). Those relative rates were then used to calculate the Pearson correlation coefficient, the ERC value, between each pair of genes.

### Enrichment Analysis of Organellar Proteomes

Pairwise ERC values between SLC30 genes and all other genes were extracted from the ERC website: https://csb.pitt.edu/erc_analysis/. Values for gene subsets of interest were used to generate visualizations and calculate enrichment statistics. Shown in **Fig 3A** are ERC values using proteomes included in **Supplementary File 1** represent key elements of mitochondrial and lysosomal proteomes that would interact with zinc transporters. Enrichment statistics were calculated using two sets of annotations to map genes to functions. First, search terms were used (show in **Fig 3B**) to search gene descriptors for each gene taken from the UCSC Genome Browser gene track. If a descriptor contained the term, the gene was mapped to that search term annotation. Also included were Mitocarta annotations (Calvo et al., 2016; Rath et al., 2021) describing mitochondrial genes. After extracting terms, enrichment was calculated in two ways. First, a one-tailed Fisher’s exact test was performed to detect an increased proportion of annotated genes with ERC values above 0.3. This test indicates that a gene being included in a particular annotation and the gene having a high ERC value are related. Second, a Wilcoxon rank-sum test was performed to look for shifts in distributions of ERC values for genes in a particular annotation compared to genes not in that annotation. AUC values, which range from 0 to 1, were calculated directly from the Wilcoxon rank-sum W statistic as W/(number of annotated genes * number of unannotated genes). Small AUC values indicate that annotated genes have *lower* ERC values than unannotated genes and large AUC values indicate that annotated genes have *higher* ERC values than unannotated genes.

### Clustering

Clusters depicted in **Fig 4A** were constructed using igraph (Gustavsen et al., 2019). Genes included were selected from genes listed in **Supplementary File 2** that represent mitochondrial complexes and ATP transporters. Lines were drawn to connect genes with ERC values greater than or equal to 0.3 and singleton genes that did not connect to other genes were removed. The optimal clustering algorithm implemented in igraph was used to define clusters.

### Protein expression and widefield microscopy

YFP-tagged C- and N-terminal in-frame fusions of human SLC30A9 sequence (NP_006336.3) were synthesized by Vectorbuilder (Chicago, IL, USA) and transfected in HeLa cells using Lipofectamine 3000 (ThermoFisher, Waltham, MA, USA). Before transfection, the cells were grown in plastic culture dishes in DMEM supplemented with 10% FBS, at 37°C, 5% CO_2_, in a humidified incubator and plated on glass coverslips at about 70% confluency. The cells were fixed in 4% formaldehyde (in phosphate-based solution, 5 min incubation) 16-24 hours post-transfection, permeabilized with 0.1% Triton X100 in phosphate-based solution, 5 min incubation, transferred to blocking buffer (1% bovine serum albumin and 1% goat serum in phosphate-based solution, 1-24 hours) and then treated with primary and secondary (fluorescent) antibodies. T0M20 antibodies (polyclonal, cat number PA5-52843) and Alexa 568-tagged secondary antibodies were from ThermoFisher. The imaging was performed using Nikon SP-1 microscope and analyzed using FIJI (Schindelin et al., 2012).

### Zinc imaging and qPCR

HeLa cells cultured as above and grown in 12-well culture plates or 35-mm Mattek dishes (MatTek Corp) were transfected with SLC30A9 esiRNA (Sigma, St Louis, MO, USA) and used 16-48 hours post-transfection. qPCR analysis was performed as before (Peña and Kiselyov, 2015), and the data were analyzed using the DDCt method. *ACTB* was used as a housekeeping gene; primers were optimized as described before (Peña and Kiselyov, 2015). For zinc-loading experiments, the cells were exposed to 100-300 μM ZnCl_2_ in DMEM/FBS for 16 hours. Next, the cells were washed, loaded with 5 μM Rhod-2,am (ThermoFisher) in a zinc-containing HEPES-based salt solution (HBSS, in mM: 140 NaCl, 5 KCl, 1 MgCl_2_, 1 CaCl_2_, 10 HEPES (pH 7.4) and 1 g/l glucose, supplemented with 100-300 μM ZnCl_2_) and washed again. The dishes were inserted into a closed, thermo-controlled (37°C) stage top incubator (Tokai Hit Co.) above the motorized stage of an inverted Nikon TiE fluorescent microscope equipped with a 60x optic (Nikon, CFI Plan Fluor, NA 1.4), a diode-pumped light engine (SPECTRA X, Lumencor). Emissions were detected using an ORCA-Flash 4.0 sCMOS camera (Hamamatsu) and excitation and emission filters were from Chroma. Zinc removal was performed by manual aspiration and gravity-fed application of nominally zinc-free HBSS. The recorded image stacks were analyzed using FIJI. Specifically, discrete regions of interest containing clearly identifiable mitochondria were outlined by hand and the dynamics of average fluorescence intensity in these images was analyzed and compared. Data are presented as mean ± standard error of the mean.

## Acknowledgements

This work was supported by the NIH grant 1R21NS111944 to KK and EA, and by NIH grant 1R01HG009299 to MC and NLC. We are grateful to Ms Livia Andrzejczuk for technical support.

## Supplementary figure legends

**Figure S1.**
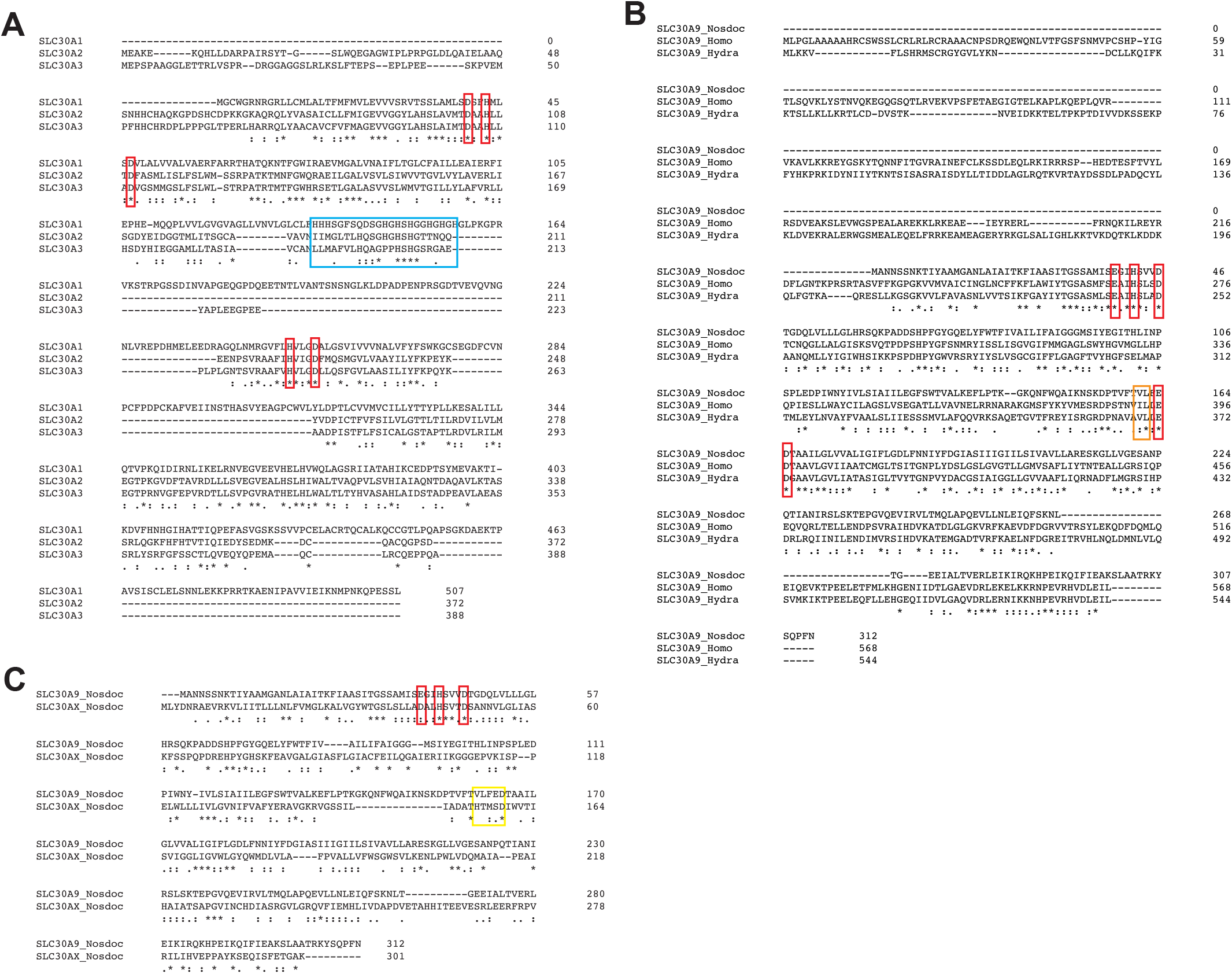
Full aminoacid alignments of human and other SLC30As including SLC30A9. Red boxes identify histidine and aspartic acid residues previously implicated in zinc permeation through SLC30As, and the aspartic and glutamic acid residues that are characteristics of SLC30A9. Blue boxes denote histidine-rich motifs. Yellow box identifies differences in the C-terminal zinc-binding motif between SLC30As and SLC30A9 in *Nostoc*.

**Figure S2.**
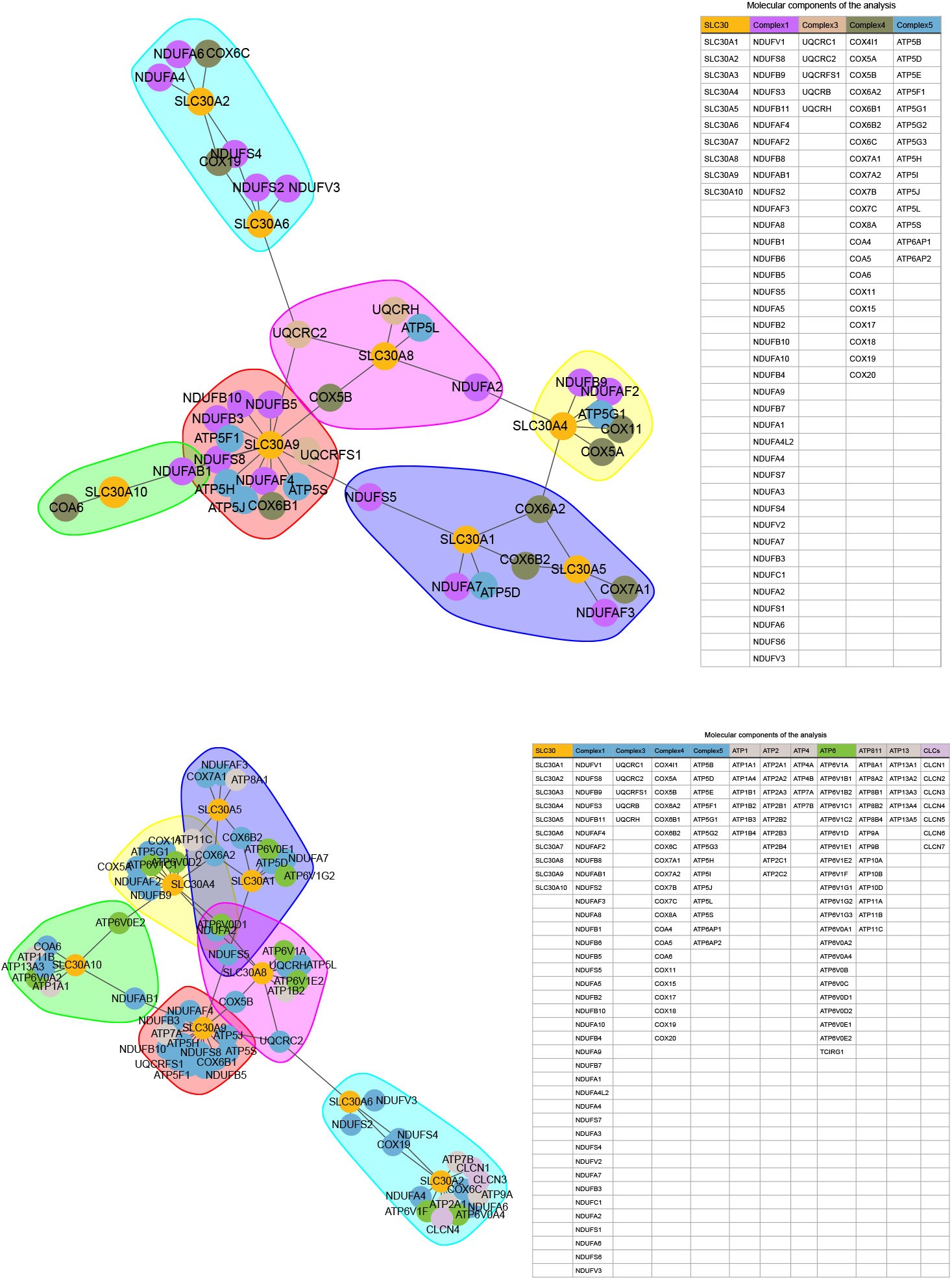
SLC30 evolutionary network clusters. Dataset from Fig 4 shown with the full list of proteins involved in the analysis.

## Notes

### Competing Interest Statement

The authors have declared no competing interest.

